# Podophyllotoxin and Quercetin *in Silico* Derivatives Docking Analysis with Cyclooxygenase-2 and Aminopeptidase-N

**DOI:** 10.1101/2021.02.26.433103

**Authors:** A.E. Manukyan, A.A. Hovhannisyan

## Abstract

The cyclooxygenase (COX) enzymes *are* tumor markers, the inhibition of which can be used in the prevention and therapy of carcinogenesis. It was found that COX-2 IS considered as targets for tumor inhibition. Aminopeptidase N (APN) is a type II membrane-bound metalloprotease associated with cancer, being identified as a cell marker on the surface of malignant myeloid cells and reached a high level of expression in progressive tumors. In anticancer therapy, plant compounds are considered that can inhibit their activity. Modeling of the COX-2 and APN enzymes was carried out on the basis of molecular models of three-dimensional structures from the PDB database [PDB ID: 5f19, 4fyq] RCSB. For docking analysis, 3D ligand models were created using MarvinSketch based on the PubChem database [CID: 5280343, 5281654]. *In silico* experiments, for the first time, revealed the possible interaction and inhibition of COX-2 and APN by quercetin and quercetin derivatives. Aspirin and Marimastat were taken to compare the results. Possible biological activities and possible side effects of the ligands have been identified.

## INTRODUCTION

Various methods and approaches are used in anticancer therapy, but the results are often not encouraging. Chemotherapy is a common treatment and does not always lead to the desired results. Anticancer drugs are used that do not have selectivity and thus cause adverse side effects. Currently, compounds of plant origin are considered in antitumor therapy, which have selective cytotoxic, cytostatic, anti-inflammatory, immunomodulatory, and other properties with low side effects.

Nevertheless, there is a constant need to develop new anticancer drugs, drug combinations and strategies of chemotherapy and photodynamic therapy through the method and scientific research of a huge number of natural drugs [Desai et al., 2008; Mukherjee et al., 2001].

In recent years, research has been rapidly developing, the use of secondary metabolites of higher plants in therapeutic agents [Gurib-Fakim, 2006; Fabricantet al., 2001]. Some metabolites, such as vincristine, vinblastine, podophyllotoxin, taxol, morphine, quercetin, etc., were isolated from plants and many of them were modified to obtain better analogs with low toxicity and / or better solubility, while maintaining high therapeutic activity. However, despite the success of this strategy, drug discoveries have only been conducted in a small number of plants [Newman, 2008]. The first step in the search for new herbal preparations or compounds was the isolation of secondary metabolites. In the past year, there have been research related to natural products and has been related to the creation of structures and stereochemistry of compounds, but in recent years there has been a large amount of research focused on their biological activity. This interdisciplinary approach is underpinned by significant progress in the development of new biological analysis methods. As a consequence, a large number of compounds isolated from plants in the past were “rediscovered” [Wagner H. et al 2012]. In 1942, Kaplan exploited the therapeutic effect of podophyllin in tumor tumors, which is still used today as an effective drug in tumor therapy. It is known that podophyllotoxin (Ptox) is usually obtained from the plants *P. hexandrum and P. peltatum*, which contain 0.2 − 4% Ptox on a dry weight basis [Chakraborty et al. 2010]. However, stocks of *Podophyllum* are limited, and root formation, where Ptox is usually stored, takes more than 5-7 years. Additional sources are plants of the family *Linacea*, genus *Linum*, which callus cultures are capable of synthesizing and accumulating lignans, in particular Ptox.

The Armenian flora contains several species that can serve as sources of Ptox [Vardapetyan et al., 2003]. The variety of biological activities and medicinal uses represented by Ptox analogs are impressive.

Batimastat is a broad-spectrum injection and metalloproteinase inhibitor (MMPI). It is the first MMPI to be clinically tested. Batimastat inhibits tumor invasion and angiogenesis, contains a thiophene heterocycle, derivatives of which are common in nature: fungi and some higher plants. Quercetin is also a well-known anti-inflammatory compound, which can be expressed on different types of cells, both in animals and in humans. Quercetin has the ability to stabilize mast cells and also has cytoprotective activity. In addition, quercetin has an immunosuppressive effect on dendritic cell function. One of the most important properties of quercetin is its ability to modulate inflammation.

The purpose and objectives of the research. The aim of this work is to study the mechanisms of action of anticancer drugs Ptox, Batimastat, Quercetin for the design of new compounds on their structural basis with an improvement/ increase in antitumor properties and a decrease in undesirable side effects.

### 1.1. Podophyllotoxin

*Linum* is a genus of annual and perennial plants of the *Linaceae* family, has medicinal properties, due to which it is widely used in medicine for the prevention and treatment of many diseases. Various medicines and additives are prepared from *Linum*. Ptox is an important plant product that is widely used in the treatment of cancer and venereal wart. Initial hopes for the possible clinical utility of Ptox as an antineoplastic agent have largely been dashed due to its side effects [Lu W et al. 2011; Van Vilet et al 2001]. Extensive structural modifications have been made to Ptox. Currently, there are various biological analogs of Ptox, which have different properties such as antimalarial, antiasmatic properties and antitumor activity [Canel C. et al. 2000, Liu YQ et al. 2005] (Figure 1).

**Figure 1.**
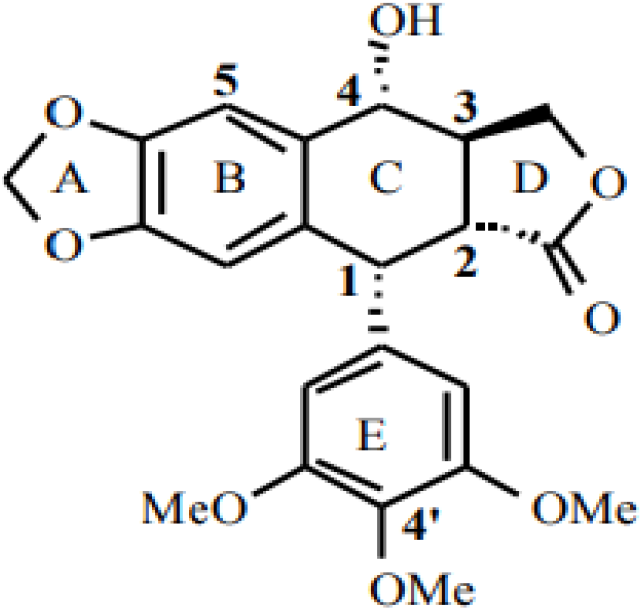
Chemical structure of the podophyllotoxin.

### 1.2. Biosynthesis of secondary metabolites in callus cultures

As is known, plant cell cultures can serve as suitable systems for studying both the biosynthesis of secondary metabolites and the metabolic pathways of their formation. The genotype of the parent plant, that is, the explant donor, significantly affects the biosynthetic potential of the cultured cells. It is assumed that cultured cells isolated from highly productive plants and tissues contain the necessary genetic information for the biosynthesis of these metabolites.

#### 1.2.1. Elicitation and biosynthesis of podophyllotoxin in callus cultures of *L. austriacum*

In vitro cultures of the genus Linum are characterized by high stability during long-term cultivation and the ability to accumulate Ptox in much larger quantities than cell cultures of *Podofillum* [Empt et al 2000, Van Uden et al1991]. The obtained undifferentiated, long-term passaged proliferating callus cultures of *L. austriacum*, grown on the MC-BN nutrient medium, were used as a model system to study the mechanisms of action of elicitors on the biosynthesis of secondary metabolites. Mannan, β-1,3-glucan, and ansymidol were used as elicitors. The investigated elicitors do not affect the growth and differentiation of callus cultures (Figure 2).

**Figure 2.**
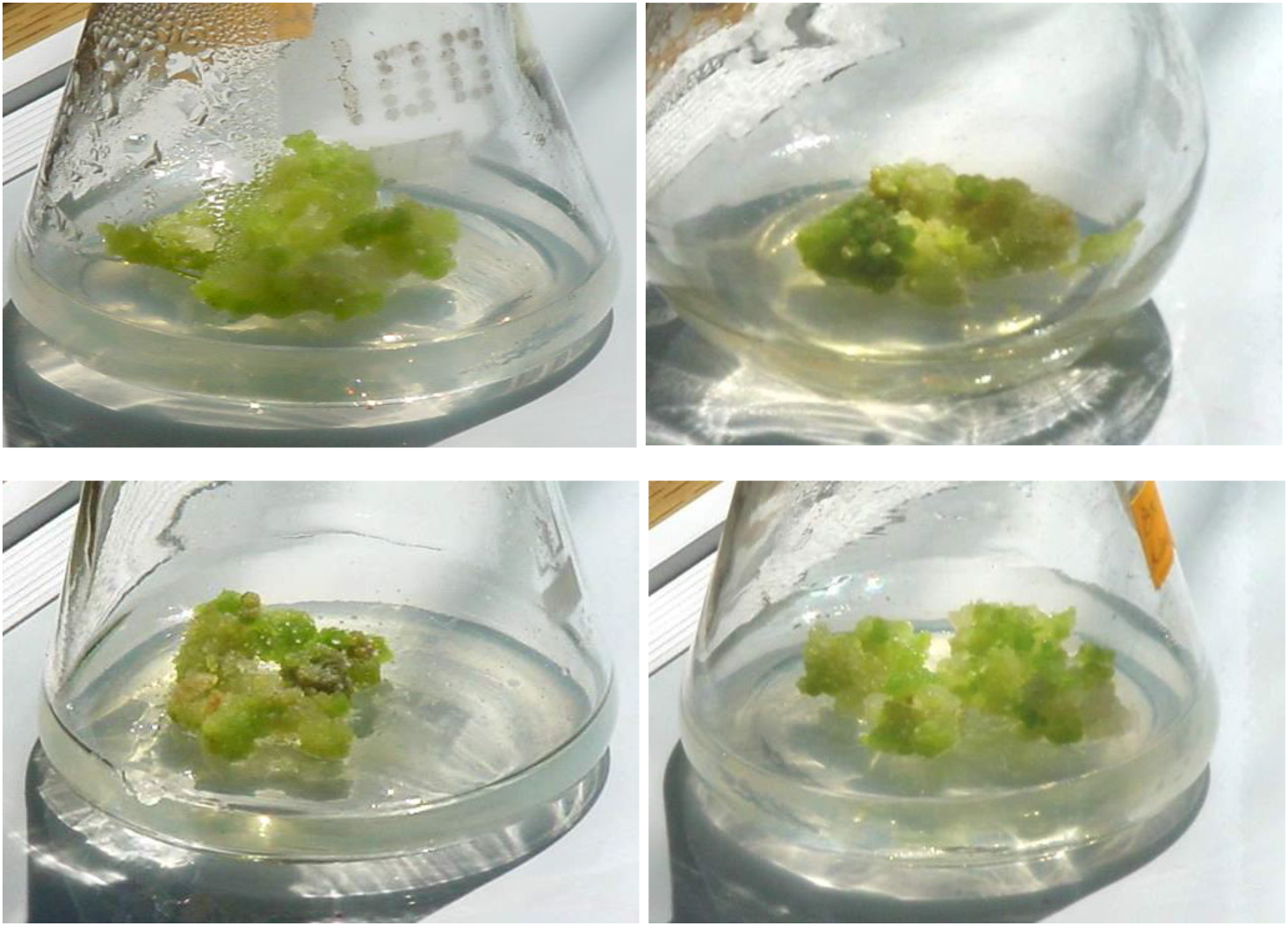
Growth of L. austriacum callus cultures on MS-BN media with various elicitors after two weeks of cultivation: a - control, b - ansymidol, c - β-1,3-glucan, d - mannan.

It follows from the results obtained that the use of elicitors of various natures strongly affects both the quantitative and qualitative spectrum of lignans synthesized in the callus cultures of *L. austriacum.*

As can be seen from the results obtained, under the influence of mannan, the biosynthesis of individual lignans in callus cultures grown in the light increases from 40% to 120%, and the total content of lignans increases from 3.87 to 7.71 mg/g dry weight (by 100%). In callus cultures grown in the dark, de-Ptox is not detected, the Ptox content decreases from 2.12 to 0.54 mg / g dry weight (by 75%), and the total lignan content decreases from 5.98 to 2.85 mg/ g (on dry weight), compared to control. The presence of Ansimidol leads to a general increase in the content of almost all lignans in callus cultures grown in the light, except for β-peltatin, which is not detected. The content of Ptox increases: from 1.28 to 1.40 mg / g dry weight for Ptox, and from 0.42 to 0.6 mg/ g dry weight for de-Ptox (about 10% and 40%, respectively), and β-peltatin from 0.82 to 1.46 mg / g dry weight (about 80%). In callus cultures grown in the dark, in the presence of Ansimidol, a high content of β-peltatin (3.65 mg / g) is observed, de-Ptox and 5-mPtox are not detected, and the content of Ptox-a decreases from 2.12 to 0.77 mg/g dry weight (or 65%), compared with control. It is noteworthy that the content of β-peltatin under the influence of β-1,3-glucan in cell cultures grown in the light increases maximally from 1.28 to 4.31 mg/ g dry weight (240% of control). At the same time, inhibition of almost all other lignans is observed. For callus cultures grown in the dark, β-1,3-glucan is an inhibitor, which may be due to inhibition of enzyme systems.

From the above, it follows that under the influence of various elicitors, selective induction of Ptox occurs in different ways. This unique property of cells and elicitors in the redistribution of secondary metabolite biosynthesis pathways can be used as a metabolic regulation tool for targeted biosynthesis of specific secondary metabolites, in particular, Ptox [Vardapetyan et al., 2003].

### 1.3. Quercetin

Quercetin, an aglycone form of plant-based flavonoid glycosides, is used as a dietary supplement and may be beneficial for a variety of diseases. Some of the beneficial effects include cardiovascular protection, antitumor, antitumor, antiulcer, antiallergic, antiviral, anti-inflammatory, antidiabetic, gastroprotective, antihypertensive, immunomodulatory, and antiinfectious [Yao Li 2016].

Due to its lipophilic nature, quercetin readily passes through cell membranes and plays a pleiotropic role in triggering various intracellular routes involved in chemoprophylaxis (eg, apoptosis, cell cycle, detoxification, antioxidant replication, cell invasion, angiogenesis). However, low bioavailability and poor solubility in water, together with rapid elimination from the body, rapid metabolism, and enzymatic degradation, significantly hinder the clinical use of quercetin as an anticancer drug (Figure 3).

**Figure 3.**
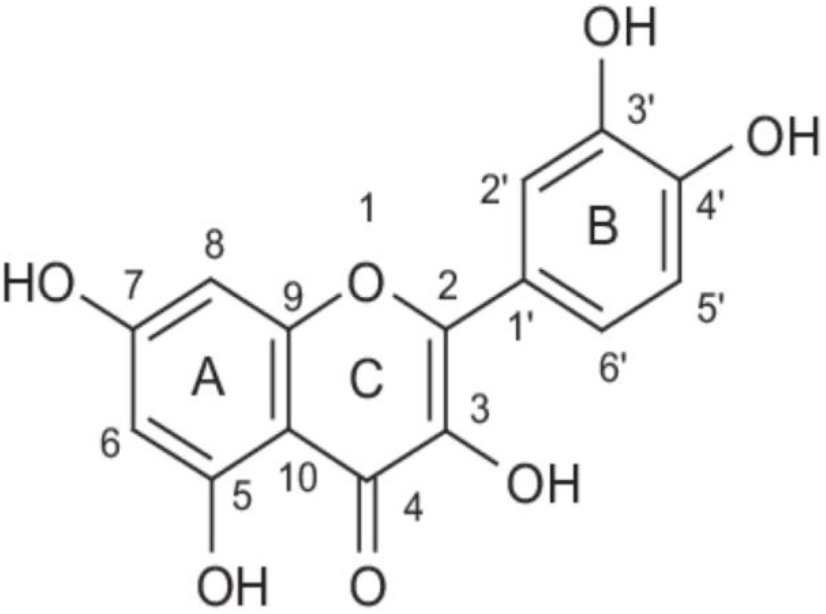
Structure of the Quercetin.

### 1.4. Low molecular weight mediators of inflammation, eicosanoids

Inflammation is a multifactorial adaptive process that occurs when the integrity of biological structures is violated. This defense mechanism is aimed at removing microorganisms, organic or inorganic material, penetrating into the tissue at the time of injury. Inflammation, regardless of etiology, develops in accordance with general pathophysiological patterns [Coussens, 2002, Yarilin 2010]. The starting point is a violation of the integrity of the cell membrane, which leads to the release of specific substances into the intercellular space, which in normal conditions are exclusively or predominantly intracellular. These are the so-called DAMPs (damage associated molecular pattern), which play the role of a biochemical “alarm” [Murakamietal. 2014; Foelletal. 2007]. DAMP interacts with specific receptors on the surface of toll-like receptors (TLRs) and in phagosomes of Nod-like receptors (NLRs) of macrophage cells, which perform the function of resident control of the immunological and structural stability of tissue [HigginsCetal. 2010; Sethietal. 2001]. Excitation of TLR and NLR triggers the synthesis of interleukins (IL) 1 and 6, tumor necrosis factor (TNFa), interferon (IFN), monocyte chemotaxis factor (CCL2) and many other biological substances that are attracted to the area of tissue damage and activate a wide range of inflammatory cells. [Fukataetal. 2009] (Figure 4). This is how the cascade of the inflammatory reaction unfolds, which, depending on the severity of the damage, goes to the systemic level with the involvement of plasma humoral factors (complement system, kinin system), changes in general metabolism and the central nervous system [Carroll 2004]. Regulation of acute inflammation is carried out according to the principle of negative feedback. In parallel with the synthesis of proinflammatory substances, the formation of IL10 molecules, resolvins (RV) and other molecules occurs, which suppress the activity and cause apoptosis of cells of the “inflammatory response”. Natural anti-inflammatory systems are connected to the process of controlling the inflammatory response [Ouyang W. et al. 2011; Schwab J. M. et al. 2007]. There is a differentiation of T-regulatory cells (CD4 + CD25 + Treg) and a special population of “alternative” macrophages (M2), blocking and destroying cells of the “inflammatory response", as well as stimulating the reparative processes of neoangiogenesis and fibrosis [Khattrietal. 2003].

**Figure 4.**
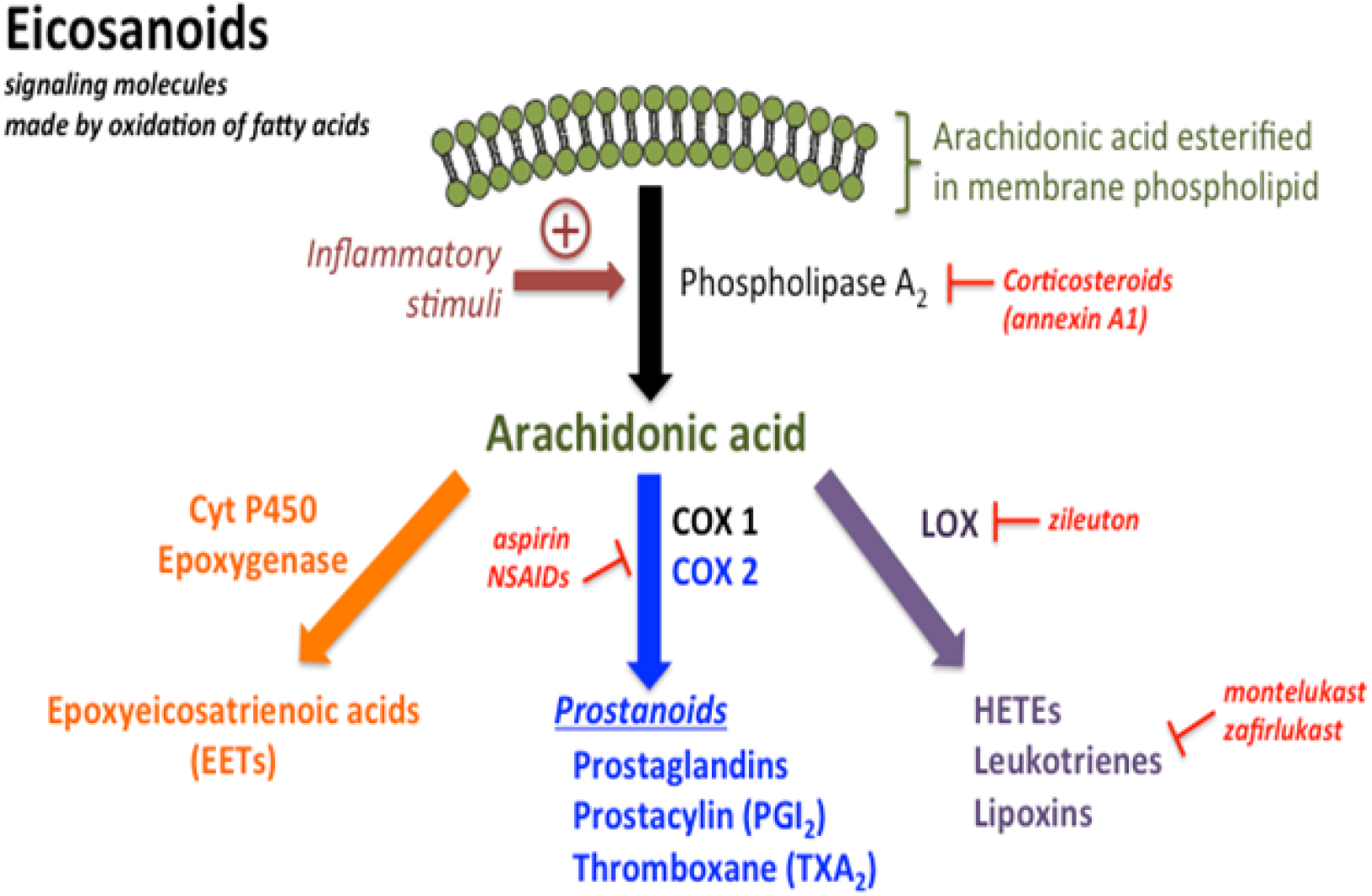
Eicosanoids synthesis.

With a favorable course of the disease or injury, against the background of the elimination of damaging factors, the concentration of DAMP rapidly decreases, IL and other anti-inflammatory substances are metabolized, and the cells of the inflammatory response lose activity and die (“cytokine deprivation”) [Ouyang et al. 2011]. Local anti-inflammatory cellular and humoral activity begins to prevail, the tissue is restored or replaced by a scar, which leads to a gradual resolution of the inflammatory response.

Molecules of various biochemical origins are involved in the regulation of the natural course of the inflammatory process. Eicosanoids are metabolites of polyunsaturated fatty acids (PFA), which occupy one of the central positions in inflammatory processes. They include prostaglandins, thromboxanes and other substances, are highly active regulators of cellular functions, have a short T1 / 2, therefore they have effects as “hormones of local action”. Excessive secretion of eicosanoids leads to a number of diseases, such as bronchial asthma and allergic reactions. Foods containing essential fatty acids are a source of PFA, but with tissue damage and inflammation, they are formed from the phospholipids of the cell membrane. The main substrate for the synthesis of human eicosanoids is arachidonic acid (Figure 4). The synthesis of arachidonic acid is carried out due to the “work” of varieties of the cytoplasmic enzyme phospholipase A2 (PLA2), which in turn is activated by pro-inflammatory signaling pathways. After the process of separating arachidonic acid from the phospholipid, it is released into the cytosol and converted into different eicosanoids depending on the cell types. The most famous is the so-called classic type of eicosanoids, which are formed due to the activity of the COX-1 and COX-2 enzymes and lipoxygenase (LOX; EC. 1.13.11.12, l) 5 and 15. The COX enzyme catalyzes the first stage of prostaglandin synthesis. The released arachidonic acid under the action of COX is converted to prostaglandin PGG2, which is reduced to prostaglandin PGH2, and that into other eicosanoids (prostaglandins, thromboxanes, prostacyclins). Prostaglandins (PGs) are the best known cellular mediators of inflammation and are derived from polyunsaturated fatty acids. Disruption of PGs biosynthesis can cause the development of severe pathological conditions: participation in the implementation of dysplastic and neoplastic processes (tumor growth), namely, in suppression of apoptotic cell death, pathological neoangiogenesis and invasion, as well as mediation of immunosuppressive functions.

#### 1.5.1. COX-2 and its biological functions

There are two main types of cyclooxygenase: COX-1 and COX-2. COX-1 is expressed in almost all mammalian tissues, and the COX-2 isozyme is constitutive in some parts of the central nervous system (hypothalamus, amygdala, hippocampus). At the same time, COX-2 is rapidly activated in response to pro-inflammatory mediators and mitogenic stimulants [Howe, et al. 2001]. Genes that encode these isozymes are characterized by different expression parameters.

One of the main cytokines that induces the corresponding cytokine-dependent signaling cascade is tumor necrosis factor (TNF), which activates pro-apoptotic receptor-mediated signaling pathways, stopping the processes of cell division and causing physiological cell death. In tumor cells, COX-2 is considered a key target in “targeted” therapy and prevention. Evidence of this conclusion is provided by the results of experimental and clinical studies, in which it was found that increased levels of COX-2 are directly related to carcinogenic processes [Dixon, et al. 2000]. A decrease in the recurrence rate of many tumors has been observed with the use of non-steroidal anti-inflammatory drugs (NSAIDs), which are COX-2 inhibitors. A fairly large number of NSAIDs are used in clinical practice. By the mechanism of inhibition of COX-2, they are subdivided into selective and non-selective, pronounced selective and non-expressed in relation to COX-1 and COX-2. But all these compounds, together with the inhibition of COX-2, also inhibit the activity of COX-1, which leads to a negative effect on the implementation of physiological processes in the body. Hence, undesirable side effects often occur when taking NSAIDs: complications from the gastric mucosa (before the appearance of gastrointestinal bleeding, ulcerative lesions), deterioration of wound healing and suppression of physiological inflammation [Stolina et al. 2000]. Due to the non-selectivity of NSAIDs, toxicity was the main reason for the development of drugs that suppress the activity of COX-2 - the so-called. “Selective inhibitors”. Thus, the search for non-toxic and effective COX-2 inhibitors continues to this day. Recently, researchers have been paying special attention to substances of natural origin that have antitumor activity and are able to selectively inhibit the activity of COX-2 [Iachininoto, et al. 2016].

#### 1.5.2. COX-2 as a promising target in anti-tumor targeted therapy

It was found that the level of COX-2 expression in tumor cells correlates with the degree of their malignancy. It was first discovered for colon tumors. In COX-2 knockout mice, an 86% reduction in the incidence of intestinal adenoma was found, and in animals heterozygous for this gene, this value was reduced by 66% compared to controls [Howeetal1999]. In experiments on female mice that repeatedly produced offspring, overexpression of COX-2 alone was sufficient to induce breast cancer in 85% of cases [Hsu et al 2000].

The regulation of COX-2 expression at the post-transcriptional level is realized through the stabilization of COX-2 transcripts. To date, it has been reliably established that the COX-2 enzyme is involved in carcinogenesis, enhancing cell proliferation (while suppressing apoptosis), pathological neoangiogenesis and cell invasion. In addition, COX-2 has a certain mutagenic effect and stimulates tumor-induced immunosuppression.

The discovery of the fact that COX-2 is directly involved in carcinogenesis has stimulated clinical trials to study the effectiveness of selective COX-2 inhibitors as agents that reduce the risk of tumors. Currently, the possibility of using COX-2 inhibitors for the prevention of tumors is being investigated. This approach has already proven its effectiveness in a model of experimental breast cancer [Alshafie, etal. 2000](Figure 5).

**Figure 5.**
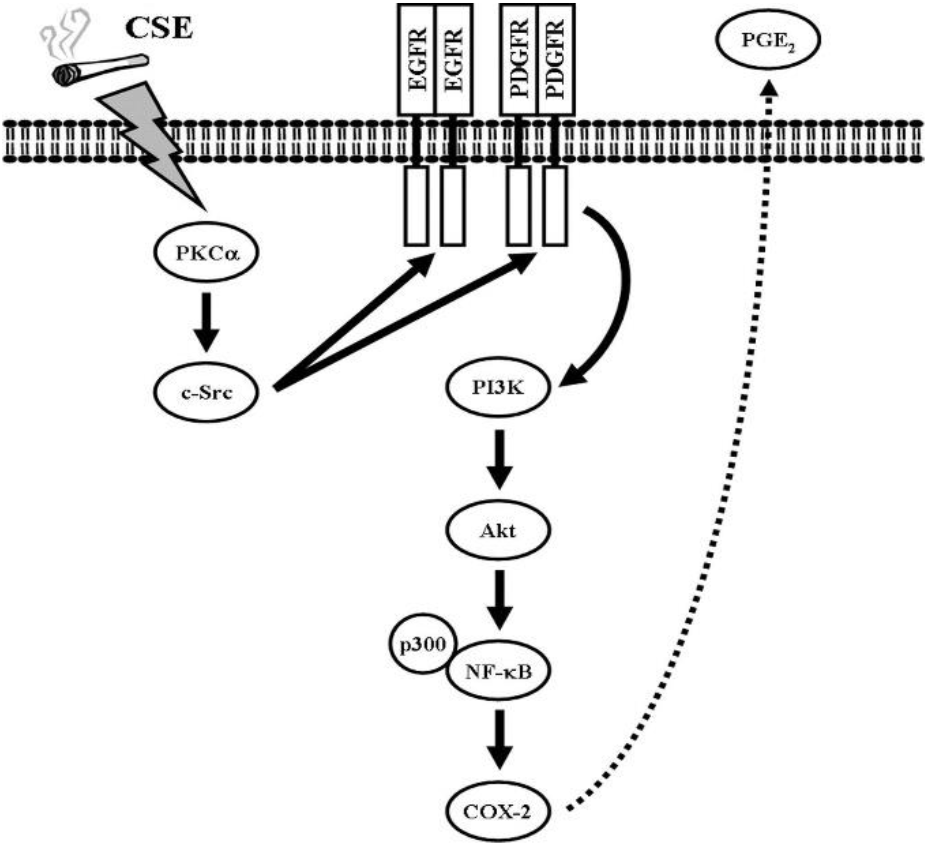
COX-2-dependent intracellular signaling cascade activated by growth factor receptors.

In conclusion, it should be noted that in recent years, natural inhibitors of COX-2 are becoming increasingly popular, as in all other cases [Zarghi, et al. 2011].

### 1.6. Metalloproteinase inhibitors

Batimastat is a broad-spectrum MMPI with activity against most major MMPs (Figure 10): interstitial collagenase (MMP-1) (IC50 3 nM), stromelysin-1 (MMP-3) (IC50 20 nM), gelatinase A (MMP −2) (IC50 4 nM), gelatinase B (MMP-9) (IC50 4 nM) and matrilisin (MMP-7) (IC50 6 nM). There is also evidence that Batimastat is a potent inhibitor of progelatinase A (MMP-14) (unpublished observations) [Knapinska, et al. 2017]. The molecule mimics the MMP substrate, so the drug is competitive. Batimastat is almost completely insoluble and therefore has a very low oral bioavailability [Corbel et al. 2001] (Figure 6). Thus, the only way to use Batimastat is to directly enter various body cavities (peritoneal Intraperitoneal injection of Batimastat produces stable plasma concentrations with an elimination half-life in humans of up to 28 days, presumably because the drug is gradually absorbed from the abdominal cavity into the bloodstream [MacDonald, et al. 1997].

**Figure 6.**
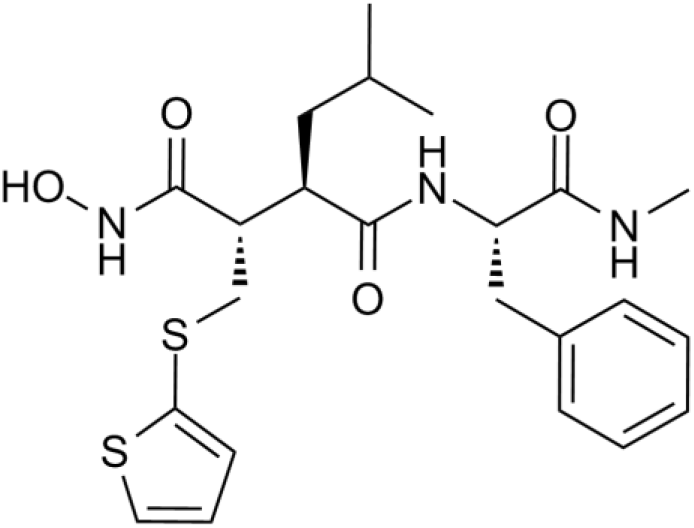
Chemical structure of the Batimastat.

#### 1.6.1 Aminopeptidase-N

Aminopeptidase N (APN, CD13, EC 3.4.11.2) is a type II membrane-bound metalloprotease [Stange et al 1996]. It is involved in the enzymatic cleavage of peptides, in endocytosis, and as a signaling molecule it regulates complex and varied processes, including cell migration, cell survival, viral uptake, and angiogenesis [Mina-Osorio et al 2009]. APN has also been associated with cancer, being identified as a cell marker on the surface of malignant myeloid cells [Shipp M.A. et al. 1993; Pasqualini R. et al. 2000] and reached a high level of expression in progressive tumors, including breast, ovarian and prostate cancer [Röcken C., et al. 2005; Di Matteo P et al. 2011]. Vascular endothelial growth factor (VEGF), a key regulator of angiogenesis, induces APN expression early in tumor growth [Bhagwat SV et al. 2001], again highlighting the role of this enzyme in angiogenesis, a process critical to the sustained growth of most tumors [Hanahan & Weinberg 2011] (Figure 7). However, APN substrates in the context of angiogenesis are still unknown. The only well-known substrate is angiotensin III in the renin-angiotensin metabolic pathway, in which APN cleaves the NH2-terminal arginine residue of angiotensin III to form angiotensin IV. APN has previously been investigated as a target for inhibiting vascularization and tumor growth [Khakoo AY et al. 2008].

**Figure 7.**
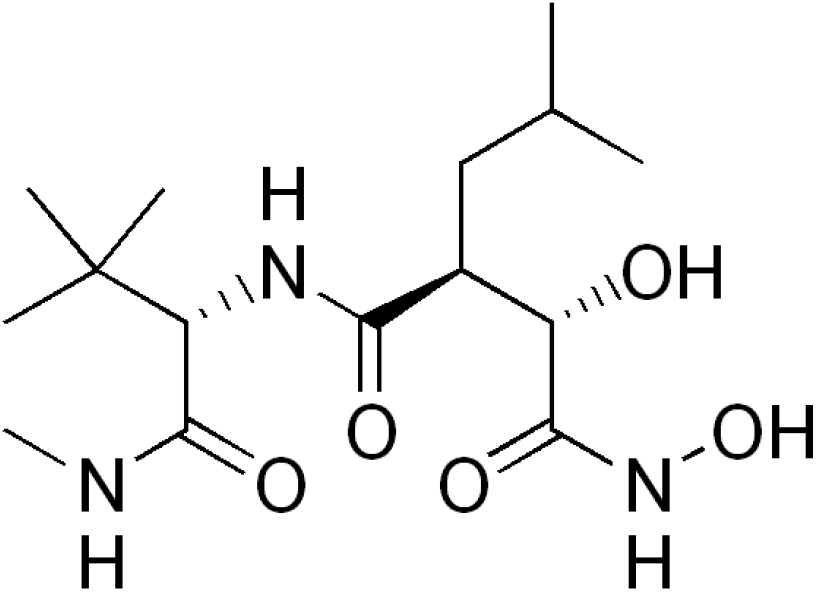
Chemical structure of the Marimastat

Tumor growth depends on a complex microenvironment in which malignant cells interact with various other types of cells: endothelial cells of the blood and lymphatic circulation, mesenchymal stromal cells, cancer-related fibroblasts, and various cells derived from bone marrow such as lymphocyte species and myeloid suppressor cells. origin [Nguyen, et al. 2009; Joyce, et al. 2009]. Some of these cell populations are involved in the induction of desmoplasia and angiogenesis. Indirect evidence suggests a role for APNs in regulating these aspects of tumor progression.

**Figure.**
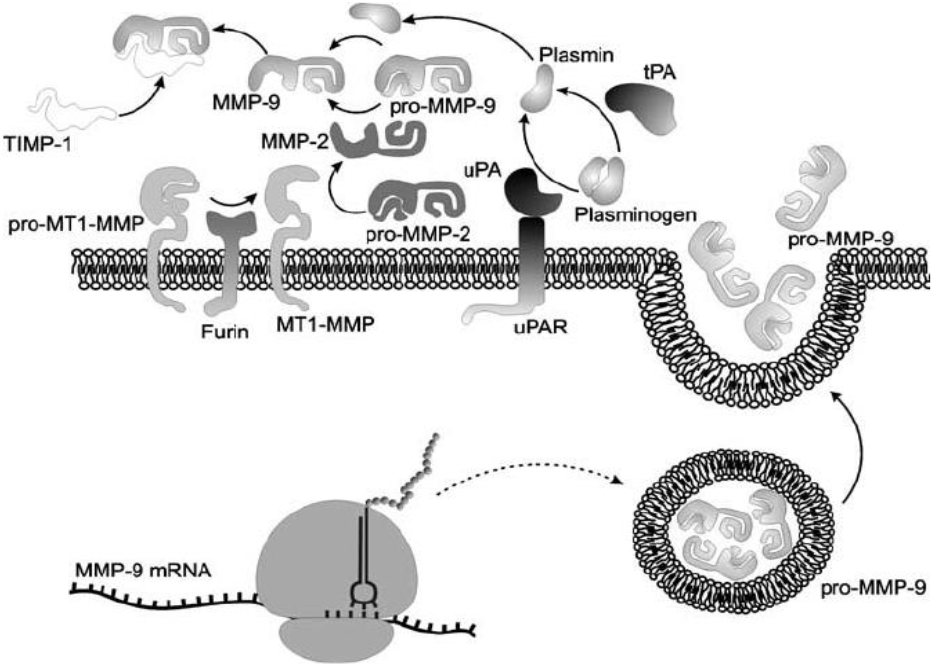
Aminopeptidase-N regulation.

Indeed, APN is expressed by both cancer cells and normal cells such as vascular endothelial cells, neutrophils and stromal cells [Di Matteo, et al. 2011], but it remains unclear where and how APN acts in the regulation of tumor angiogenesis.

## Material and methods

### 2.1. Obtaining callus cultures of L. austriacum

Callus cultures of *L. austriacum* were obtained from sterile seedlings. The seeds were sterilized in 3% and 5% hydrogen peroxide for 40 minutes, then washed with sterile distilled water. The seeds were germinated in the dark in twice diluted MS medium [Murashige & Skoog 1962]. For callusogenesis, after 7 days, the resulting *L.austriacum* seedlings were damaged and transferred into 100 ml Erlenmeyer flasks containing 25 ml of sterilized modified MS medium (MS-BN-medium or MS-N) with the following additives: 30 g/ L sucrose, 8 g/ L agar-agar, 0.1 mg/ L thiamine-HCl, 100 mg/ L myo-inositol, 1 mg/ L nicotinic acid, 1 mg/ L pyridoxine-HCl, phytohormones (0.5 mg/ L BAP and 0.005 mg/ L α- NAA or 0.4 mg/ L α-NAA) (all firms “Sigma”, USA), pH was adjusted to 5.8 with 0.1 M KOH, pre-autoclaved under 1 atmosphere pressure (Steam Sterilizer, Israel).

Obtained callus (Fig. 8) were subcultured at 26 ° ± 1 ° C, relative humidity 70%, under continuous illumination on agar media MS-BN and MS-N. The original callus cultures retained their regenerative properties and biosynthetic potencies during the entire research period.

**Figure 8.**
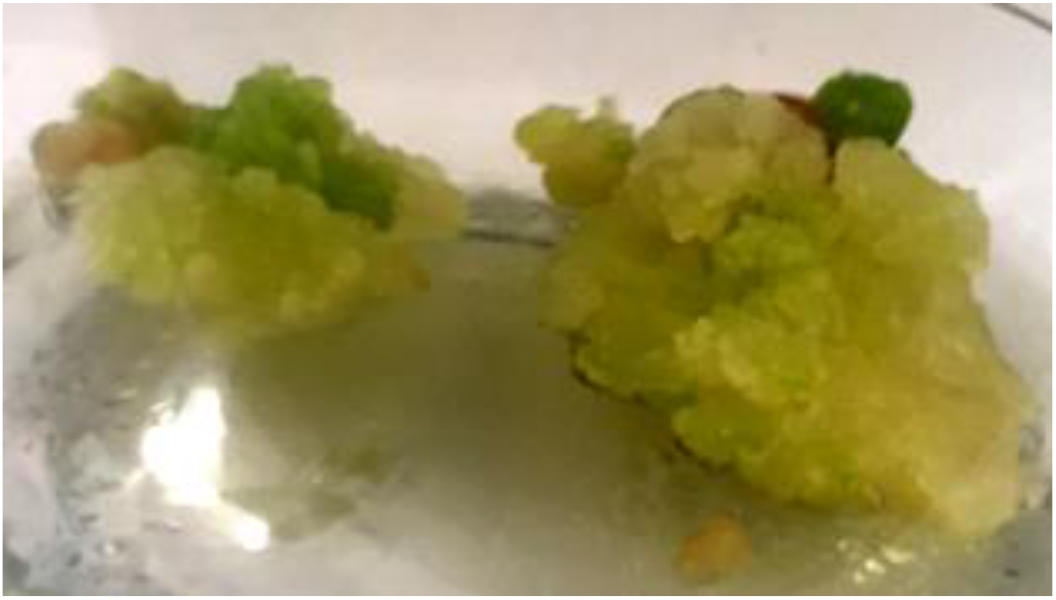
Callus culture of *L.austriacum*.

### 2.2 Extraction of podophyllotoxin and its derivatives

Sampling for chemical analysis was carried out from freeze-dried callus tissue of L. austriacum. Tissues (0.2 g each) were ground in a porcelain mortar, mixed with 2 ml of ethanol, and homogenized 2 times for 30 seconds on a high-speed homogenizer MPW-302 (Mechanika Precysyjna, Poland) with continuous cooling. 6 ml of distilled water was added to the resulting suspension and adjusted to pH 5.4 with 0.6% o-phosphoric acid, after which 0.1 ml of distilled water containing 0.1 mg of β-glucosidase (Sigma, USA). The mixture was incubated for 1 hour in a water bath at 35 ° C. The resulting mixture was added with 12 ml of ethanol to improve the dissolution of lignans and incubated for 10 min at 70 ° C in an ultra-incubator. Then the suspension was centrifuged for 15 min at 12000 g, the supernatant was separated and used for chromatographic analysis [Vardapetyan et al. 2002]. The prepared preparations were stored at −20 ° C.

### 2.3 Analysis of podophyllotoxines by HPLC

Determination of the relative content of Ptox, 5-mPtox, de-Ptox, 〈- and ®-peltatins in L. austriacum cell cultures was carried out using HPLC on an HPLC-Termo Quest device (Termo Quest, Germany) using Spherisorb ODS columns −2 (Sigma, USA), in a gradient system: water containing 0.1 ml / L of 90% o-phosphoric acid (A) and methanol-acetonitrile (B) in a 1: 1 ratio. Chromatographic conditions: 0-4 min - in 5% B, the concentration of which was linearly increased to 100% within 40 min. Elution rate 1 ml / min, sample volume 25 μl. Commercial preparations of Ptox, 5-mPtox, de-Ptox (Roth, Germany) were used as standards. All peaks corresponding to the values of Rt - Ptox, 5-mPtox, de-Ptox, peltatins were automatically subjected to spectral analysis by comparing them with the absorption spectra of commercial preparations using a special program. The concentration of Ptox-s was determined by the equation: C = √S × √350 / ɛ where: S-peak area, ɛ - extinction equal to 29000. The number of Ptox-s was measured automatically based on the area of each peak at 290 nm.

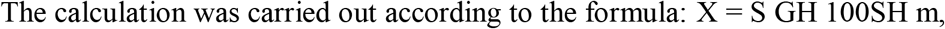

-where S is the area of the peak of Ptox in the sample, S is the area of the peak of Ptox in the standard solution, m is the weight of the dry plant in g, X is the% content of Ptox in the dry plant, GH is the weight of the Ptox in the weight of the standard solution, about 0.025 g. A 10% methanol solution of Ptox-a (0.0026 mg/ml) was used as a standard [Vardapetyan et al. 2003].

### 2.4 Target screening Batimastat and Ptox

The search for targets for Batimastat and Ptox was carried out on the basis of verified articles from the Pubmed database. The chemical structures of Batimastat and Ptox were taken from the PubChem chemical structure bank [CID: 10607, 5362422] [Bolton, et al. 2008]. Derivatives of the latter were constructed using the chemical editor MarvinSketch. In order to optimize the ligands, the degrees of freedom were calculated, polar hydrogen atoms were added, and the atomic charges were calculated. Based on the chemical structures of the ligands, using TargetNet, possible targets for each ligand were determined [Zhi-Jiang Yao et al. 2016]. From ~ 5000 targets, duplicates were removed and targets belonging to *H. sapiens* were selected (Table 3). Docking analysis was performed for each target and the 2 with the highest minimum binding energy were selected. For Batimastat - APN, for Ptox - COX-2.

Since the structure and functions of the two ligands were well studied, *in silico* models of compounds derived from Batimastat and Ptox were constructed on their basis.

Current research is aimed at identifying a potential link between COX-2 expression and ABCG2 [Sorokin 2004]. Expression of COX-2 may be associated with the MDR phenotype through the overexpression of some ABC transporters.

### 2.5. Preparation of chemical structures of ligands

To create a library of chemical compounds, the structures of podophyllotoxin and batimastat were obtained from the PubChem database, *in silico* modified derivatives of these compounds were used as derivatives, which were constructed using MarvinSketch. Aspirin and Marimastat were studied as a comparison of the results obtained. The resulting compound structures were verified based on Lipinski’s rule of five. In order to optimize the ligands, the degrees of freedom were calculated, polar hydrogen atoms were added, and the atomic charges were calculated.

### 2.6. Preparation of protein molecules

Crystallographic data on protein structure belonging to Homo sapiens were taken from the PDB RCSB database (PDB: 5F19, 4FYQ) (COX-2, APN) [Berman, et al. 2002]. The crystallographic structure of COX-2 was in a complex with aspirin, for this reason it was decided to remove aspirin and repair the structure of COX-2, which is acetylated in the position of Ser530 when aspirin interacts. Structural changes were carried out using PyMOL [DeLano, 2002]. After that, the binding sites were checked using the MOLE 2.5 program. As a result, two major binding pockets were identified in proteins (Figure 9).

**Figure 9.**
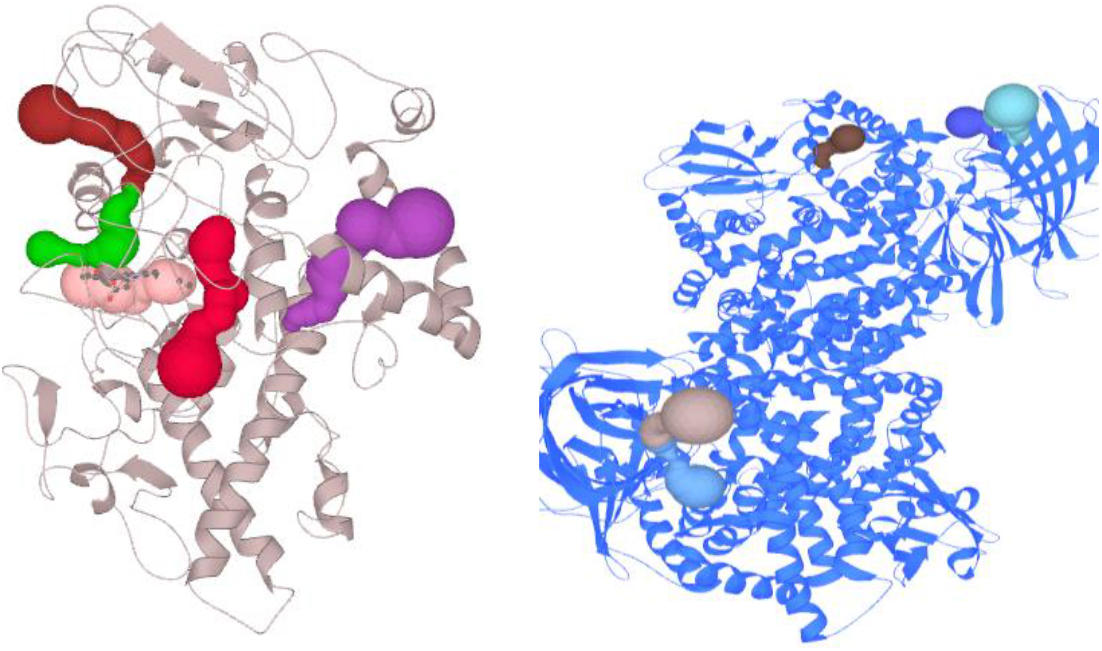
Binding pockets for COX-2, APN: 4 main binding pockets.

At the initial stage, the spatial and energy indicators of the interaction of the studied compounds with the active centers of proteins were determined. The basic parameters for COX-2 were a virtual box with dimensions 28X28X30, for APN- 34X28X34. The search depth was 8, 16, 32, 64, 128, 256, 512, after which 10 compounds were selected, docking analysis of which showed the highest binding energy with proteins. For visualize the docking results, the PyMOL program [DeLano L. 2002] was used, which is a free cross-platform molecular graphics system.

### 2.6. Analysis of the similarity of in silico research methods with in vitro methods

The calculation of physicochemical parameters, as well as the prediction of ADMET parameters (Absorption, Distribution, Metabolism, Excretion and Toxicity) were carried out using the SwissADME programs and PASS online, which are intended for use in the design of medicinal compounds. The PASS online toxicity prediction includes an assessment of the probability of presence (Pa) and inactivity (Pi) with values ranging from 0 to 1, as well as an assessment of activity and toxicity on human cell lines.

## RESULTS AND DISCUSSION

### 3.1. Biosynthesis of lignans in callus cultures of *L. austriacum* L

Ptox and its derivatives have become the subject of research for their unique properties (antibacterial, antiviral, antioxidant and antitumor). As is known, plant cell cultures can serve as suitable systems for studying both the biosynthesis of secondary metabolites and the metabolic pathways of their formation. The genotype of the parent plant, that is, the explant donor, significantly affects the biosynthetic potential of the cultured cells. It is assumed that cultured cells isolated from highly productive plants and tissues contain the necessary genetic information for the biosynthesis of these metabolites. In this case, it is also necessary to take into account the preservation of the biosynthetic potencies of plants during the transition to cell cultures. Thus, although the roots of *P. peltatum* and *P. hexandrum* have the highest Ptox content [Jackson & Dewick 1984; Fay & Ziegler 1985], but callus cultures of these plants grow very poorly and cannot serve as stable sources of Ptox [Moraes-Cerdeira et al. 1998; Nadeem et al. 2000]. The obtained cell cultures of L. austriacum were cultured on agar medium MC containing α-NAA (MC-N medium) and BAP + α-NAA (MC-BN medium), which are considered optimal for cell cultures of the genus *Linum* [Petersen & Alfermann 2001], on light.

It should be noted that in cultures grown in the dark on MS-N medium, the total content of lignans increases from 1.92 to 2.65 mg / g dry weight (by 38%), and the accumulation of biomass significantly decreases compared to cultures passaged under continuous illumination, where the content of α-peltatin and Ptox increases from 0.49 to 0.71 mg/g dry weight (by 46%) and from 0.36 to 0.46 mg/g dry weight (by 27%), respectively (Fig. 16). It follows from the results that L. austriacum cell cultures grown on the MC-BN nutrient medium synthesize approximately twice as many lignans as those grown on the MC-N medium. The MS-BN medium is optimal both for the growth of callus cultures and for the biosynthesis of secondary metabolites. The experiments showed that the MS-BN medium is the most optimal medium for the growth of the biomass of L. austriacum callus cultures. Cell cultures of L. austriacum on MS-BN medium accumulate 5-mPtox, Ptox, ®-peltatin as the main product.

As previously shown, the content of lignans in callus cultures can be almost doubled by using various elicitors as instruments for metabolic regulation of targeted biosynthesis of specific secondary metabolites, in particular, Ptox [Vardapetyanetal. 2003]. Thus, the obtained callus cultures can be used as producers of podphyllotoxin.

### 3.2. Screening and selection of suitable targets

The chemical structures of the ligands: Ptox, Batimastat, Quercetin were taken from the PubChem bank [CID: 10607, 5362422, 5280343] [Bolton, et al. 2008]. Based on the structures of Ptox-a, Batimastat, all possible targets for each ligand were determined using TargetNet. From ~ 5000 targets, repeats were removed and targets belonging to *H. sapiens* were selected (Table 1) and taking into account their potential proto-oncogenic properties. Thus, we have chosen the following targets:

**Table 1.**

| Nº | Ligand | COX-2 | APN | ABCG2 |
| --- | --- | --- | --- | --- |
| 1. | **44** | VAL 434,<br>TYR385,<br>LEU503,<br>TYR 355,<br>ARG120 | D439,<br>D473,<br>A474,<br>H388 | ILE409,<br>VAL546,<br>ASN436<br>PHE439, |
| 2. | **1** | VAL434,<br>TYR385,<br>LEU503,<br>TYR 355,<br>ARG120 | D439,<br>D473,<br>A474,<br>H388 | ILE409,<br>VAL546,<br>ASN436<br>PHE439,<br>THR435 |
| 3. | **12** | VAL434, | D439, | ILE409, |
|  |  | TYR385,<br>LEU503,<br>TYR 355,<br>ARG120 | D473,<br>A474,<br>H388 | VAL546,<br>ASN436<br>PHE439,<br>THR435 |
| 4. | **43** | VAL<br>434, TYR385,<br>LEU 503, TYR<br>355,<br>ARG120 | D439,<br>D473,<br>A474,<br>H388 | ILE409, VAL546,<br>ILE409,<br>VAL546,<br>ASN436<br>PHE439, |
| 5. | **4** | VAL<br>434, TYR385,<br>LEU 503, TYR<br>355,<br>ARG120 | D439,<br>D473,<br>A474,<br>H388 | ILE409,<br>VAL546,<br>ASN436<br>PHE439, |
| 6. | **5** | VAL<br>434, TYR385,<br>LEU 503, TYR<br>355,<br>ARG120 | D439,<br>D473,<br>A474,<br>H388 | ILE409,<br>VAL546,<br>ASN436<br>PHE439, |
| 7. | **14** | VAL<br>434, TYR385,<br>LEU 503, TYR<br>355,<br>ARG120 | D439,<br>D473,<br>A474,<br>H388 | ILE409,<br>VAL546,<br>ASN436<br>PHE439, |
| 8. | **21** | VAL 434,<br>TYR385,<br>LEU503, | D439,<br>D473,<br>A474, | ILE409,<br>VAL546,<br>ASN436 |
|  |  | TYR 355,<br>ARG120 | H388 | PHE439, |
| 9. | **16** | VAL 434,<br>TYR385,<br>LEU503,<br>TYR 355,<br>ARG120 | D439,<br>D473,<br>H388 | ILE409,<br>VAL546,<br>ASN436<br>PHE439,<br>THR435 |
| 10 | **22** | VAL 434,<br>TYR385,<br>LEU503,<br>TYR 355,<br>ARG120 | D439,<br>D473,<br>A474,<br>H388 | ILE409,<br>VAL546,<br>ASN 436,<br>PHE 439,<br>THR435 |

Neoplasia marker - COX-2 (EC 1.14.99.1) is involved in carcinogenesis, enhancing cell proliferation (while suppressing apoptosis), pathological neoangiogenesis and cell invasion and immunosuppression of cancer cells, the inhibition of which can be used in the prevention and therapy of carcinogenesis [Uddin et al, 2016]. Since COX-2 was inhibited in the database, acetyl-ser530COX-2 (inhibited form of COX-2 by aspirin at ser530) was taken. With the help of PyMOL, acetyl-ser530 was replaced by ser530.

Aminopeptidase-N (APN / CD13, EC 3.4.11.2) is a membrane-bound Zn-dependent type II MMP, it carries out enzymatic N-terminal cleavage of amino acid residues of peptides, participates in endocytosis and, as a signaling molecule, participates in the regulation of complex and varied processes, including cell migration, cell survival, receptor mediated viral endocytosis, and angiogenesis. APN is considered as a target for inhibiting tumor vascularization and growth. Mammalian APN, or CD13, is a tumor marker that is overexpressed on the cell surface of almost all major tumor forms [Saiki I. et al. 2016].

### 3.3. Docking analysis results

As noted above, the identification of the active components of plant extracts, new phytopharmaceutical compounds and their mechanisms of action is an urgent task. Secondary plant metabolites can be used as drugs in the treatment of many diseases. Lignans, in particular Ptox, are classified as herbal antitumor agents. Interest in Ptox and its derivatives is mainly due to their pronounced cytotoxic and antiviral activity [de Bruin et al. 2014; Gordaliza et al. 2000].

Batimastat is a broad spectrum injectable drug and inhibitor of MMPs. It is a competitive peptidomimetic and the first MMPI to be clinically tested. Batimastat inhibits tumor invasion and angiogenesis, but exhibits adverse side effects [Bauvois & Dauzonne 2006].

Quercetin is one of the important bioflavonoids present in over twenty plant materials and is known for its anti-inflammatory, anti-tumor, etc. Inhibits the activity of COX-2.

For a detailed study of the interaction of derivatives with the receptor, a docking analysis was performed to identify possible binding sites for each ligand with each protein. The data obtained indicate the possible binding of all ligands to targets. From the data obtained, 10 compounds were selected taking into account the thermodynamic parameters [Table 1].

When arachidonic acid interacts with COX-2, an electron is transferred to heme from Y385 with the formation of a tyrosyl radical. The nonspecific inhibitor, aspirin, acetylates S530, which blocks the entry of the substrate (arachidonic acid) into the active site. Docking analysis of Ptox- a, Batimastat and quercetin, as well as derivatives with the COX-2-aspirin complex, was performed. From the results it was revealed that with the modified amino acid S530, “competitive” binding with COX-2 occurs.

As a comparison, docking analysis was performed with specific protein inhibitors: aspirin, marimastat.

The results showed that COX-2 binds to Aspirin with an energy of −6.5 kcal/Mol, which is a relatively average interaction result. Docking analysis of COX-2 in combination with aspirin revealed that all ligands interact with amino acids VAL 434, TYR 385, ARG 120, HIS 351. In the case of APN, the results showed that Marimastat binds with an energy of −8.5 kcal/Mol forming hydrogen bonds, as well as electrostatic and hydrophobic interactions with amino acids ALA353, ARG935, ASP473, ALA474.

The results of docking analysis showed that Ptox probably interacts with COX-2 with positively charged NH/ N groups HIS351 with hydrogen/ electrostatic, and hydrophobic interactions with amino acids THR383, VAL447. Batimastat probably interacts with the amino acid residues COX-2, which are located close to the active site - LIS446, THR383, GLY533, LIS467, mainly by hydrophobic interactions. Quercetin interacts with a.k.o. COX-2 GLU465, THR383, VAL538, probably with electrostatic, hydrogen and hydrophobic interactions, respectively. The derivatives bind to a.k.o THR383, GLU465, VAL538, which are close to the active center of COX-2, and binds by hydrogen and electrostatic interactions. The results of docking analysis showed that Batimastat binds to the amino acid residues of COX-2, mainly by hydrophobic interactions, while Ptox and derivatives form not only hydrophobic, but also hydrogen bonds and electrostatic interactions (Figure 10).

**Figure 10.**
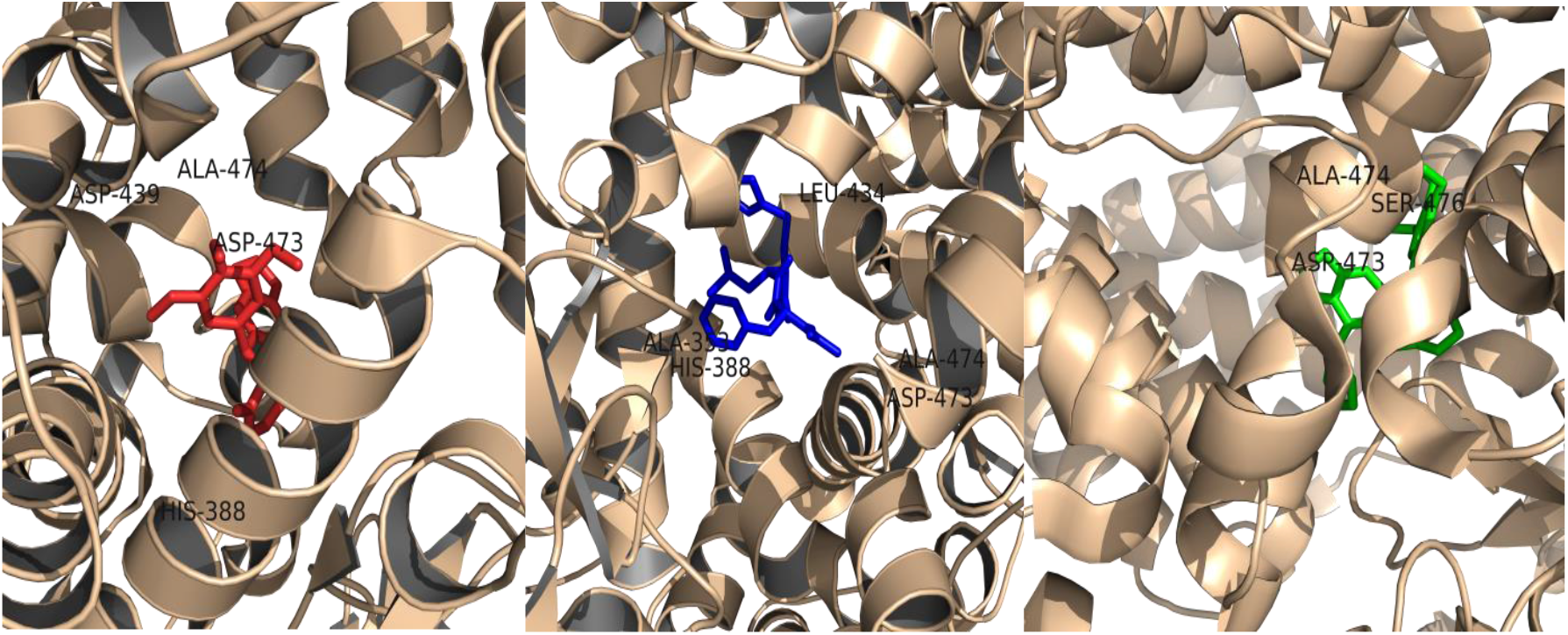
Interaction with APN: a) Ptox, b) Batimastat,c) Quercetin.

In the case of APN, the results showed that Ptox interacts with an energy of −7.8 interacts with a.k.o ASP439, ASP473, ALA474, HIS388. Batimastat with an energy of −9.1 with a.k.o A353, H388, L434, D473, A474, forming hydrogen bonds, as well as electrostatic and hydrophobic interactions. Quercetin interacts with an energy of −6.9 with a.k.o ALA474, ASP473, SER476. Batimastat with an energy of −7.5 s a.k.o ILE409, VAL546, PHE439, forming hydrogen bonds, as well as electrostatic and hydrophobic interactions. Quercetin interacts with −7 a.k.o ILE409, VAL546, ASN436, PHE439.

Docking analysis of the derivatives showed that all ligands interact with proteins. Were selected 10 ligands, taking into account the thermodynamic parameters. Interactions of ligands with a.k.o. shown in Table 1.

It is noteworthy that the results of docking analysis of Batimastat and Ptox showed no interaction with APN at position S476, given that the minimum binding energy of *in silico* derivatives with APN increases by 8% compared to other ligands. Therefore, it can be assumed that the binding of derivatives with S476 provides greater activity and stability of this complex.

### 3.4. SwissADME results

Based on the chemical structural formulas for each ligand, using Cell Line Cytotoxicity Predictor (CLC-Pred), the cytotoxic action of chemical compounds in nontransformed and cancer cell lines was predicted in silico. The results in table 2.

**Table 2.**
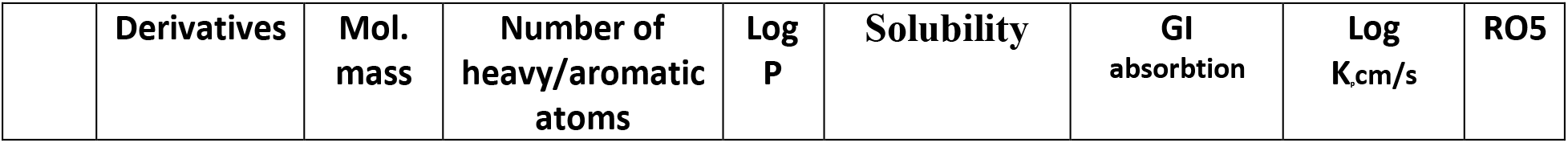
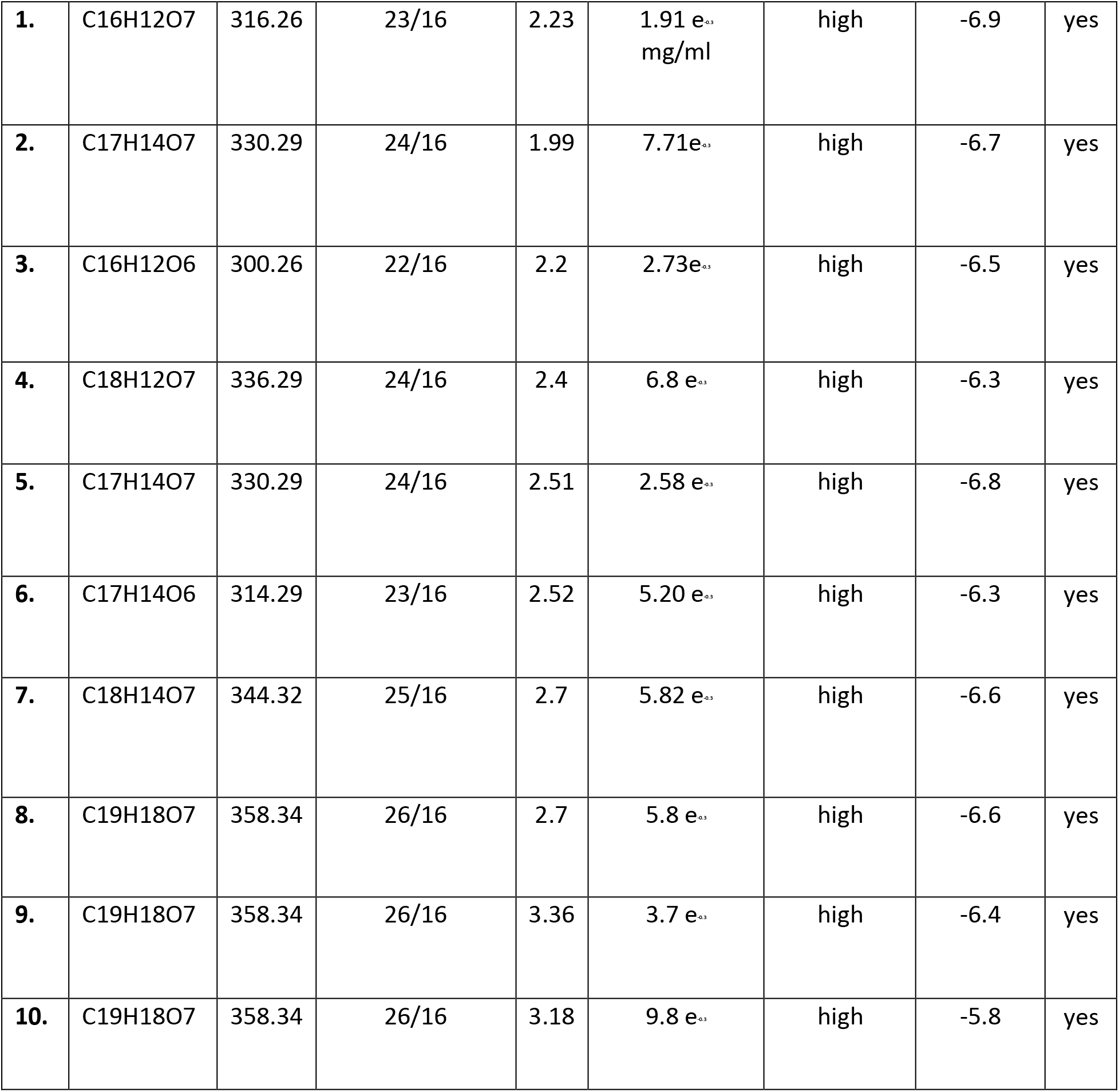
Physico-chemical parametrs of the derivatives.

## Conclusion

The drug discovery and development process aims to provide affordable drugs that are the safest, most effective and specific to improve longevity, quality of life. However, this process is very complex, time consuming and resource intensive, as well as multidisciplinary experience and innovative approaches. According to the latest estimates, plant extracts are the basis of pharmaceutical innovation and the most optimal method for identifying new biologically active components.

In this work, we studied the production of callus cultures of Linum austriacum, which are capable of synthesizing and accumulating lignans, in particular Ptox. Ptox has strong cytotoxic properties. Previously, extensive structural modifications to Ptox have been undertaken, which have led to the clinical introduction of two semi-synthetic analogs, etoposide and teniposide. Batimastat inhibits tumor invasion and angiogenesis and contains a thiophene heterocycle, derivatives of which are widespread in nature: fungi and some higher plants. Based on the chemical structures of the anticarcinogenic compounds Ptox, Batimastat, Quercetin, we have designed compounds with a potential improvement / increase in antitumor properties, with a decrease in unwanted side effects. As a result of *in silico* experiments, the possible interaction of Ptox, Batimastat, Quercetin and their in silico derivatives with neoplasia markers - cyclooxygenase-2-COX-2 and with Zn-containing type II metalloproteinase - APN and breast cancer resistance protein - ABCG2 - was revealed for the first time. The obtained results were compared with the results of interaction of specific inhibitors with receptors, and a “competitive” binding of the ligands Ptox, Batimastat, Quercetin and in silico derivatives with the inhibited form of COX-2, which was in a complex with Aspirin, was revealed. The physicochemical parameters of Ptox, Batimastat, Quercetin and in silico derivatives were calculated and the probable activities on tumor cell lines, as well as side effects on tissues.

Author: Manukyan Amalya

Institute: Russian-Armenian University, Bioengineering, Bioinformatics and Molecular Biology Street: Hovsep Emin 123 str.

City: Yerevan

Country: Armenia

## Notes

### Competing Interest Statement

The authors have declared no competing interest.

## REFERENCES

1. Alshafie G. A. et al. Chemotherapeutic evaluation of Celecoxib, a cyclooxygenase-2 inhibitor, in a rat mammary tumor model //Oncology reports. – 2000. – V. 7. – №. 6. – P. 1377–1382.

2. Andjelic B. et al. A single institution experience on 314 newly diagnosed advanced Hodgkin lymphoma patients: the role of ABVD in daily practice //European journal of haematology. – 2014. – V. 93. – №. 5. – P. 392–399.

3. Arakelyan V., Babayan Y., Potikyan G. Determination of constant rates of adsorption of ligand on DNA: analysis of correlation functions //Journal of Biomolecular Structure and Dynamics. – 2000. – V. 18. – №. 2. – P. 231–235.

4. Bennett R. N., Wallsgrove R. M. Secondary metabolites in plant defence mechanisms //New phytologist. – 1994. – V. 127. – №. 4. – P. 617–633.

5. Berman H. M. et al. The protein data bank //Acta Crystallographica Section D: Biological Crystallography. – 2002. – V. 58. – №. 6. – P. 899–907.

6. Bhagwat S. V. et al. CD13/APN is activated by angiogenic signals and is essential for capillary tube formation //Blood. – 2001. – V. 97. – №. 3. – P. 652–659.

7. Bolton E. E. et al. PubChem: integrated platform of small molecules and biological activities //Annual reports in computational chemistry. – 2008. – V. 4. – P. 217–241.

8. Bourboulia D., Stetler-Stevenson W. G. Matrix metalloproteinases (MMPs) and tissue inhibitors of metalloproteinases (TIMPs): Positive and negative regulators in tumor cell adhesion //Seminars in cancer biology. – Academic Press, 2010. – V. 20. – №. 3. – P. 161–168.

9. Bramhall S. R. et al.Marimastat as first-line therapy for patients with unresectable pancreatic cancer: a randomized trial //Journal of Clinical Oncology. – 2001. – V. 19. – №. 15. – P. 3447–3455.

10. Bramhall S. R. et al.Marimastat as maintenance therapy for patients with advanced gastric cancer: a randomised trial //British journal of cancer. – 2002. – V. 86. – №. 12. – P. 1864–1870.

11. Brill S. J. et al. Need for DNA topoisomerase activity as a swivel for DNA replication for transcription of ribosomal RNA. – 1987.

12. Cao Y. et al. Intracellular unesterified arachidonic acid signals apoptosis //Proceedings of the National Academy of Sciences. – 2000. – V. 97. – №. 21. – P. 11280–11285.

13. Carroll M. C. The complement system in regulation of adaptive immunity //Nature immunology. – 2004. – V. 5. – №. 10. – P. 981–986.

14. Chakraborty A. et al.An efficient protocol for in vitro regeneration of Podophyllum hexandrum, a critically endangered medicinal plant. – 2010.

15. Corbel M. et al.Inhibition of bleomycin-induced pulmonary fibrosis in mice by the matrix metalloproteinase inhibitor batimastat //The Journal of pathology. – 2001. – V. 193. – №. 4. – P.538–545.

16. Cos P. et al. Anti-infective potential of natural products: how to develop a stronger in vitro ‘proof-of-concept’ //Journal of ethnopharmacology. – 2006. – V. 106. – №. 3. – P. 290–302.

17. Coussens L. M., Werb Z. Inflammation and cancer //Nature. – 2002. – V. 420. – №. 6917. – P.860–867.

18. Croteau R., Kutchan T. M., Lewis N. G. Natural products (secondary metabolites) //Biochemistry and molecular biology of plants. – 2000. – V. 24. – P.1250–1319.

19. Csizmadia F. JChem: Java applets and modules supporting chemical database handling from web browsers //Journal of Chemical Information and Computer Sciences. – 2000. – V. 40. – №. 2. – P.323–324.

20. Cunha I. et al. Factors that influence the yield and composition of Brazilian propolis extracts //Journal of the Brazilian Chemical Society. – 2004. – V. 15. – №. 6. – P. 964–970.

21. Cybulski G. R. et al. Luque rod stabilization for metastatic disease of the spine //Surgical neurology. – 1987. – V. 28. – №. 4. – P. 277–283.

22. Daina A., Michielin O., Zoete V. SwissADME: a free web tool to evaluate pharmacokinetics, drug-likeness and medicinal chemistry friendliness of small molecules //Scientific Reports. – 2017. – V. 7. – P.42717.

23. DeLano W. L. The PyMOL Molecular Graphics System. De-Lano Scientific, San Carlos, CA, USA //http://www.pymol.org. – 2002.

24. Desai A. G. et al. Medicinal plants and cancer chemoprevention //Current drug metabolism. – 2008. – V. 9. – №. 7. – P. 581–591.

25. Ding X. Z., Tong W. G., Adrian T. E. Blockade of cyclooxygenase-2 inhibits proliferation and induces apoptosis in human pancreatic cancer cells //Anticancer research. – 1999. – V. 20. – №. 4. – P. 2625–2631.

26. Dixon D. A. et al. Post-transcriptional control of cyclooxygenase-2 gene expression The role of the 3′-untranslated region //Journal of Biological Chemistry. – 2000. – V. 275. – №. 16. – P. 11750–11757.

27. Dixon J. et al. Expression of aminopeptidase-n (CD 13) in normal tissues and malignant neoplasms of epithelial and lymphoid origin //Journal of clinical pathology. – 1994. – V. 47. – №. 1. – P. 43–47.

28. Eccles S. A. et al.Control of Lymphatic and Hematogenous Metastasis of a Rat Mammary Carcinoma by the Matrix Metalloproteinas Inhibitor Batimastat (BB-94) //Cancer research. – 1996. – V. 56. – №. 12. – P.2815–2822.

29. Fabricant D. S., Farnsworth N. R. The value of plants used in traditional medicine for drug discovery //Environmental health perspectives. – 2001. – V. 109. – №. Suppl 1. – P. 69.

30. Fauré M. et al. Antioxidant activities of lignans and flavonoids //Phytochemistry. – 1990. – V. 29. – №. 12. – P.3773–3775.

31. Filimonov D. A. et al. Prediction of the biological activity spectra of organic compounds using the PASS online web resource //Chemistry of Heterocyclic Compounds. – 2014. – V. 50. – №. 3. – P. 444–457.

32. Filleur F. et al. Antiproliferative, anti-aromatase, anti-17β-HSD and antioxidant activities of lignans isolated from Myristica argentea //Planta medica. – 2001. – V. 67. – №. 08. – P.700–704.

33. Flex A. et al. Human cord blood endothelial progenitors promote post-ischemic angiogenesis in immunocompetent mouse model //Thrombosis research. – 2016. – V. 141. – P.106–111.

